# Exposure of mammary cells to lipid activates gene expression changes associated with ER-negative breast cancer via chromatin remodeling

**DOI:** 10.1101/2020.12.13.422540

**Authors:** Shivangi Yadav, Ranya Virk, Carolina H Chung, David Van Derway, Duojiao Chen, Kirsten Burdett, Hongyu Gao, Zexian Zeng, Manish Ranjan, Gannon Cottone, Xiaoling Xuei, Sriram Chandrasekaran, Vadim Backman, Robert Chatterton, Seema Ahsan Khan, Susan E Clare

## Abstract

Improved understanding of local breast biology that favors the development of estrogen receptor negative (ER-) breast cancer (BC) would foster better prevention strategies. We have previously shown that overexpression of specific lipid metabolism genes is associated with the development of ER- BC. We now report results of exposure of MCF-10A cells and mammary organoids to representative medium- and long-chain polyunsaturated fatty acids. This exposure caused a dynamic and profound change in gene expression, accompanied by changes in chromatin packing density, chromatin accessibility and histone posttranslational modifications (PTMs). We identified 38 metabolic reactions that showed significantly increased activity, including reactions related to one-carbon metabolism. Among these reactions are those that produce S-adenosyl-L-methionine for histone PTMs. Utilizing both an *in-vitro* model and samples from women at high risk for ER- BC, we show that lipid exposure engenders gene expression, signaling pathway activation, and histone marks associated with the development of ER- BC.

## Introduction

Breast cancer is a heterogeneous disease with different molecular subtypes that are characterized, at a minimum, by the expression of the estrogen receptor (ER), progesterone receptor (PR) and Human epidermal growth factor receptor 2 (HER2)/neu (*1*). Although multiple statistical tools have been developed to quantify breast cancer risk (*2*), they do not predict breast cancer subtypes. Current breast cancer prevention with selective estrogen receptor modulators (SERM) and aromatase inhibitors decreases the risk of estrogen-receptor (ER) positive breast cancer sub-types, but not those without ER expression (*3-5*). Thus, determining the etiologic/biologic factors that favor the development of ER-negative breast cancer will potentially enable the development of both strategies to identify women at risk for ER-negative disease as well as targeted preventive and therapeutic agents.

Given the poor understanding of the genesis of sporadic ER-negative breast cancer, we set out to study this using the contralateral, unaffected breast of patients with unilateral breast cancer as a model. Studies of metachronous contralateral breast cancer show a similarity in the ER status of the contralateral cancer to the index primary *(6-8)*. Therefore, the contralateral unaffected breast (CUB) of women undergoing surgical therapy for newly diagnosed unilateral breast cancer can be employed as a model to discover potential markers of subtype-specific risk. In a previous study, we performed Illumina expression arrays on epithelial cells from the CUB of breast cancer patients, and identified a lipid metabolism (LiMe) gene signature which was enriched in the CUBs of women with ER-breast cancer (*9*). Among these are genes that control critical steps in lipid and energy metabolism. We validated this signature in an independent set of 36 human samples and re-confirmed the above results in fresh frozen tissues obtained from a new set of ER+ and ER-breast cancer patients, each time using laser capture microdissection (LCM) to obtain epithelial cells from tumor and CUB samples (*10*). Again, we found significantly higher expression of LiMe genes in CUBs from women with ER-breast cancer, compared to both CUBS from women with ER+ breast cancer, and breast epithelium from a control group of women undergoing reduction mammoplasty. However, the specific genes comprising this overexpressed set had no specific function or group of functions in common and did not suggest specific mechanistic explanations as to why lipid metabolism pathways would aid ER-breast cancer development. In the present study, we address possible mechanistic explanations for our previous observations.

Cellular metabolism is a complex sequence of reactions in response to a cell’s microenvironment that have profound effects on cellular function (*11*). Major reprogramming of cellular energetics is one of two emerging hallmarks of cancer (*11*). Metabolic re-wiring is required to provide the energy required to enable continuous growth and proliferation of the cancer cells. The past century has witnessed intensive investigation of metabolic pathways in cancer, in particular that of aerobic glycolysis commonly called as the Warburg effect (*12*). However, this is not the singular anomaly in the metabolically altered cancer cell. In addition to glucose and glutamine, fatty acids are an extremely important energy source (*13*). Altered lipid metabolism is posited to be a driver of carcinogenesis in various cancers, including ovarian (*14*), prostate (*15, 16*), liver (*17*) and triple negative breast cancer (*18, 19*). Increased lipid metabolism has also been shown to serve as a survival signal that enables tumor recurrence and has been suggested as an Achilles heel for combating breast cancer progression (*20*). Despite this recognition of the importance of fatty acid metabolism, its role of in the transformation of a normal cell to the malignant state is largely unknown. Metabolomic studies of the concentrations of several free fatty acids in primary breast tumors, including linoleate, palmitate, and oleate, as a function of breast cancer subtype have revealed significant differences across the subtypes, with the highest concentrations in basal-like breast cancer (*21*). Conjugation of long-chain fatty acids to carnitine for transport into the mitochondria and subsequent fatty acid oxidation (FAO) was observed to be highest in basal-like breast cancers, followed by luminal B ∼HER2-enriched, with luminal A tumors displaying the lowest levels (*21*). Another study, which utilized Raman spectroscopy to interrogate tissue, revealed that histologically normal breast tissue centimeters removed from the breast malignancy have significantly higher polyunsaturated fatty acid levels compared with normal tissue from cancer-free subjects (*22*).

Metabolites from intermediate metabolism are the substrates used to generate chromatin modifications, underlining a complex relationship between metabolism and epigenetics. Key to the crosstalk between metabolism and chromatin structure, is that the kinetic and thermodynamic properties of the chromatin modification reactions are commensurate with the dynamic range of the physiological concentrations of the corresponding intermediates in metabolism (*23*). For example, the substrates for histone methylation and acetylation reactions often have cellular concentrations that are commensurate with enzyme Km values, and thus are sensitive and responsive to changes in metabolism. Historically, glucose-derived carbon has been considered the primary source of acetyl-coA for histone acetylation. In the nucleus, acetate may be a minor source. Recently, however, data from McDonnell and colleagues has revealed that lipids reprogram metabolism to become a major carbon source for histone acetylation (*24*). This reprogramming was shown to have significant effects on gene expression. Therefore, we sought to determine if the LiMe signature we observed in the CUBs of ER-patients is associated with chromatin modifications and histone PTMs secondary to changes in metabolism fostered by exposure to medium and long chain fatty acids.

## Results

### Lipid facilitates transcriptional reprogramming in non-transformed mammary cells

We established an *in vitro* model by exposing estrogen and progesterone receptor (PR) negative MCF10A cells to octanoate, a medium chain eight-carbon fatty acid. Due to its small size and lipophilic nature octanoate does not depend on fatty acid transport proteins to traverse cell membranes and is readily oxidized in the mitochondria to form acetyl-CoA (*25, 26*). We performed RNA-seq to determine the effects of octanoate treatment on gene expression in the MCF10A cells. RNA-seq analysis revealed that 24 hours of octanoate treatment produces a transcriptional profile that is completely distinct from vehicle-treated controls (**Fig. 1A, Fig. S1A-B**). Genes with initially low expression (negative values of ln(*E*_*ctrl*_/*E*_*ctrl,avg*_)) are upregulated (corresponding to positive values of ln(*E*_*oct*_/*E*_*ctrl*_)) while genes with initially high expression (positive values of ln(*E*_*ctrl*_/*E*_*ctrl,avg*_)) are downregulated upon octanoate treatment (corresponding to negative values of ln(*E*_*oct*_/*E*_*ctrl*_))(*27*). More specifically, there is a clear trend for initially highly expressed genes in the control condition to be downregulated upon octanoate treatment while genes with initial low expression in the control condition were upregulated. Differential expression analysis revealed a total of 2132 upregulated and 632 downregulated genes (FDR=0.01) in the octanoate treated cells (**Supplementary Fig. 1C**). Pathway enrichment analysis of the differentially expressed genes induced by the 5mM octanoate treatment was performed and the top 25 upregulated and downregulated pathways are shown in Fig. 1B. Specifically, this analysis revealed that among the top altered biological processes are second messenger mediated signaling, the Notch signaling pathway, adenylate cyclase-activating adrenergic receptor signaling, cell morphogenesis and differentiation. In contrast, downregulated genes are involved in cell cycle processes, transcriptional regulation of tumor suppressor genes such as p53, and cell cycle checkpoints (**Fig. 1B**). Additional gene set enrichment analysis (GSEA) investigating top pathways with coordinated upregulation or downregulation of genes demonstrated that the top pathways associated with octanoate treatment included positive regulation of cell morphogenesis, a process involved in differentiation, as well as several oncogenic pathways associated with breast tumorigenesis, including *ERBB, WNT*, and *NOTCH* signaling pathways (**Fig. 1C**). Subsequent leading-edge analysis of these top upregulated signaling pathways-Lipid storage pathways (I), Wnt pathway (II), Notch signaling (III) and ERBB pathway (IV) shows clear association of core enrichment genes with octanoate treatment across replicates (**Fig. 1D**). Network analysis of octanoate-associated pathways identified by GSEA analysis revealed linked clusters involved with the nervous system and a second, separate group of linked clusters involved with growth factor stimulation, regulation of the *MAPK* cascade, and *ERBB* signaling (**Fig. 1E**). Finally, using real-time qPCR we validated the expression of a number of genes that GSEA analysis determined were significantly upregulated with octanoate treatment (**Fig. 1F**). Thus, treatment with medium chain fatty acids induces significant changes in transcription (*28*).

**Fig. 1.**
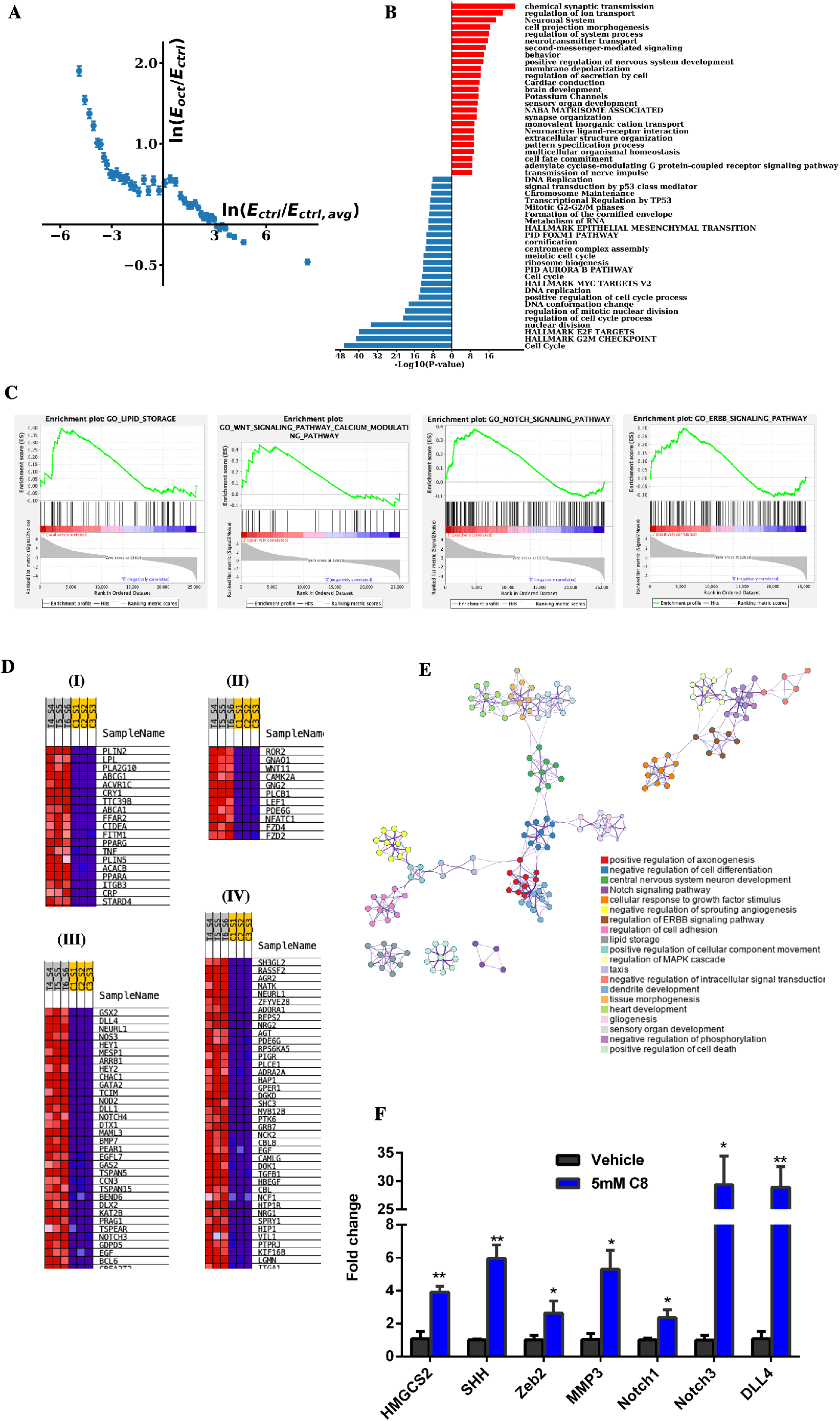
Lipid-rich environment enables transcriptional reprogramming in mammary epithelial cells. (A) 24-hour treatment of MCF10A cells with 5mM octanoate results in a completely distinct transcriptional profile compared to untreated controls. *E*_*ctrl*_ is the expression of genes in the control condition across all 3 control replicates, *E*_*ctrl,avg*_ is the average expression for the control condition across all genes and replicates, *E*_*oct*_ is the expression of genes across all 3 octanoate replicates. *E*_*ctrl*_/*E*_*ctrl,avg*_ represents the ratio of expression of a particular gene to the average expression across all control cells. Thus, a positive value of 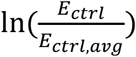 corresponds to genes that are highly expressed in the control conditions while a negative value of 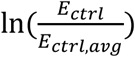 corresponds to genes that have an initial lower expression in the control condition. *E*_*oct*_/*E*_*ctrl*_ represents the ratio of expression of a particular gene for octanoate-treated versus vehicle control-treated cells. Genes with initially low expression are upregulated while genes with initially high expression are downregulated upon octanoate treatment. (B) Gene ontology analysis of differentially expressed genes induced by octanoate treatment. Upregulated and downregulated genes were first identified using DESeq2 (FDR < 0.01, |logFC| = 2) for 5mM octanoate treated cells compared to vehicle-treated control cells. Pathway enrichment analysis was performed on identified differentially expressed genes with annotations from online pathway databases (KEGG, Hallmark, Canonical Pathways, Reactome, BioCarta) and Gene Ontology Biological Processes. Pathway enrichment was ranked by p-value on a −Log_10_ scale and a selection from the top 25 pathways associated with upregulated genes (in red) and downregulated genes (in blue) are shown.(C) GSEA analysis of Gene Ontology Biological Processes showing top pathways associated with octanoate treatment with FDR < 0.1 related to differentiation, cell signaling, and metabolic processes. (D) List of core enrichment genes differentially expressed in treated replicates -T4, T5, T6 versus control replicates-C1, C2, C3 (I) Lipid storage pathways (II) Wnt pathway (III) Notch pathway (IV) ERBB pathway each pathway as identified by GSEA leading edge analysis. (E) Network analysis of pathways associated with the octanoate phenotype in GSEA analysis of Gene Ontology Biological Processes. (F) qPCR analysis of genes associated with the NOTCH pathway. Two genes, NOTCH3 and DLL4 show significant upregulation upon 5mM octanoate treatment compared to other identified genes such as NOTCH1. Statistical significance was determined by the unpaired t-test with Welch’s correction.

### Evaluating the lipid composition of the serum of ER- and ER+ BC patients

Next, we investigated whether dietary lipids, which are mainly long chain fatty acids (LCFAs), have a similar effect on the gene transcriptional profile to that of MCF10A cells. In order to determine the specific lipid(s) to evaluate experimentally, we sought to determine the differences in the percent composition of lipid species as a function of ER expression in serum from patients who had donated CUB samples for our original studies (*9, 10*). A comprehensive lipid profile of these serum samples was performed by the Northwest Metabolic Research Center at University of Washington, with measurement of more than 700 lipids. For each of the measurements, the association between the measured value and ER status was evaluated using regression models, adjusting for BMI, age, and menopausal status. ER was a categorical variable used to describe subjects having ER + or ER – cancers, or controls undergoing reduction mammoplasty. As the purpose of this experiment was to identify a lipid for ensuing experiments, lipid species were ranked for effect size comparing serum from patients subjects with ER-disease to those with ER+ disease. Three of the top four lipid species with the largest effect size were noted to contain linoleic acid: cholesterol ester (CE) 18:2, phosphatidyl choline (PC)16:0/18:2 and triacylglycerol (TAG) 54:6-FA18:2. Linoleic acid as a free fatty acid ranked 11^th^ in the analysis. Linoleic acid is the most highly consumed polyunsaturated fatty acid in the human diet (*29*), its presence in serum CE has been strongly correlated with intake (*30*), and its concentration in adult adipose tissue has more than doubled in the past half century (*31*). Therefore, all subsequent studies were carried out using linoleic acid (LA).

### Linoleic acid influences chromatin packing behavior

The state of chromatin is intimately linked with the regulation of gene transcription, undergoing dynamic changes between transcriptionally active and inactive states. Thus, our next step was to explore the changes in chromatin structure of fatty acid treated MCF10A cells by employing partial wave spectroscopic (PWS) microscopy, which quantifies chromatin packing scaling (*D*) in live cells (*32*). *D* represents the power-law scaling relationship between the 1-D size of the chromatin polymer i.e. the number of nucleotides and the 3-D space the chromatin polymer occupies. Recent evidence indicates that higher chromatin packing scaling is associated with increased intercellular and intra-network transcriptional heterogeneity as well as increased malignancy and chemoresistance in cancer cells (*27, 33, 34*). PWS was used to evaluate the effect of LA on chromatin packing scaling in live MCF10A cells. Images were obtained every 6 hours over a 24-hour period. Our results showed significant increases in chromatin packing scaling upon exposure to LA, in a manner similar to that of octanoate, suggesting that there is an increase in the dynamic range of gene expression and transcriptional gene network heterogeneity following lipid treatment (**Fig. 2A-B**). Thus, LA treatment results in changes in chromatin packing structure which are associated with a more malignant phenotype. Such significant changes in chromatin packing behavior also indicate significant changes in chromatin accessibility, which is directly associated with chromatin structure (*35*).

**Fig. 2.**
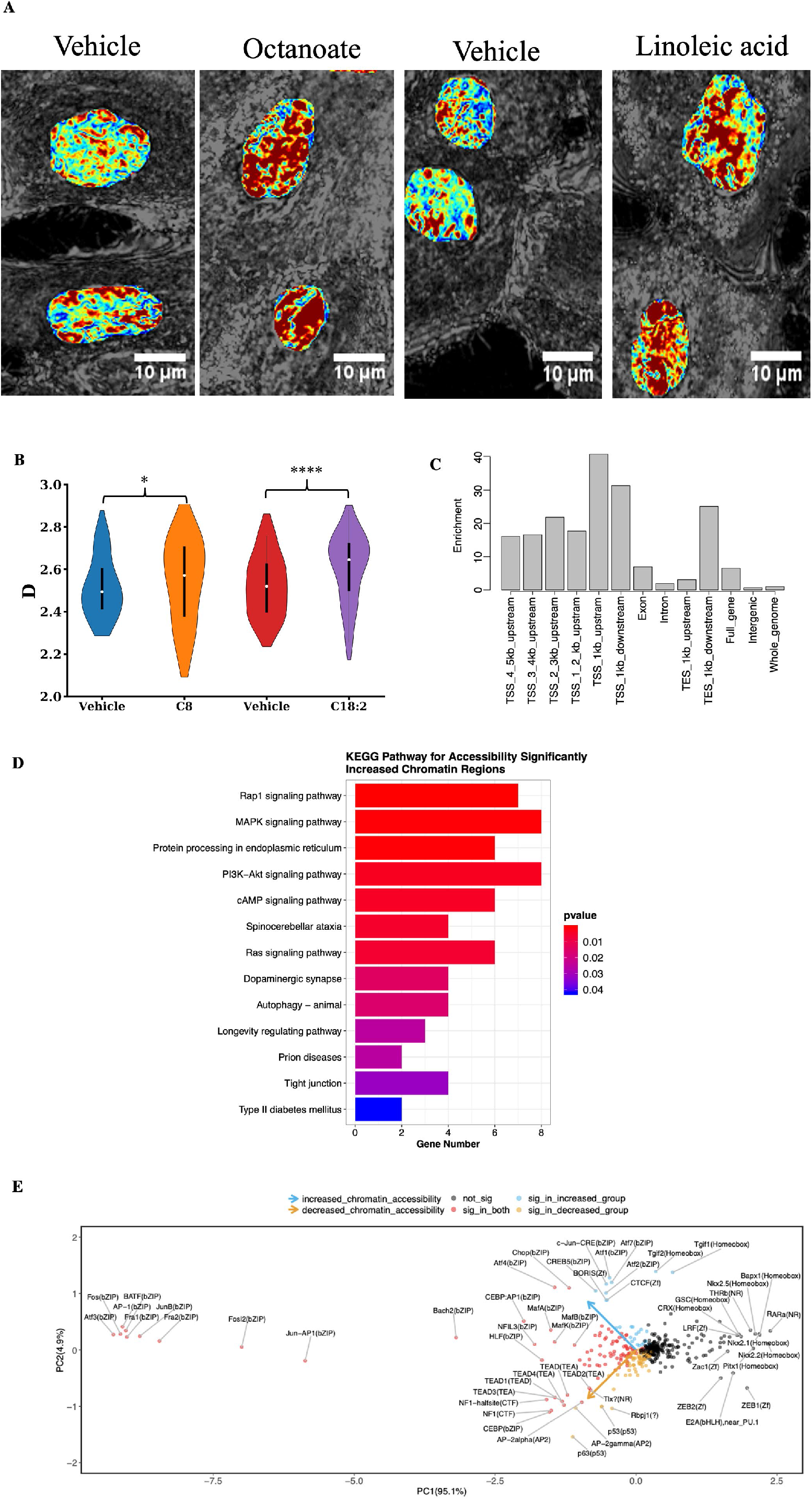
Linoleic acid alters large-scale chromatin packing behavior in MCF-10A cells. (A) Representative PWS microscopy images of MCF-10A cell nuclei at 24 hours after treatment with vehicle controls and lipids – octanoate and linoleic acid. Scale bars, 10μm. Chromatin packing scaling *(D)* map of nuclei shows an increase in chromatin packing scaling upon lipid treatment as demonstrated by an increase in red regions. (B) Changes in average chromatin packing scaling among MCF-10A cells upon treatment with vehicle controls and lipids compared to untreated cells. Significance was determined using unpaired Kolmogorov-Smirnov *t*-test (\*\*\*\**P* < 0.0001, \**P* < 0.05). Bar graphs show the mean change in intranuclear *D* across cell populations for N = 88 cells PBS (vehicle for octanoate), N = 110 cells Octanoate (C8), N = 103 cells BSA (vehicle for linoleic acid), and N = 94 Linoleic acid (C18:2). (C) Enrichment of genomic locations for 1704 open chromatin regions (FDR < 0.05, logFC > 1) in LA treated MCF-10A cells. The enrichment of peaks in each type of genomic region relative to the whole genome is shown on the y-axis. Two ATAC-seq libraries were used for the analysis. (D) Pathway analysis for the regions with increased chromatin accessibility in linoleic acid treated cells identified using the KEGG database. (E) Biplot showing changes in chromatin accessibility for specific regions identified by HOMER analysis. Motifs with a significant increase in the chromatin accessibility are shown in blue and those with a significant decrease in accessibility are shown in yellow (FDR<0.05 and |logFC| > 1).

### ATAC Sequencing reveals increased chromatin accessibility in regulatory regions of genes in the MAPK and cAMP signaling pathways in lipid treated mammary cells

To acquire more detailed insight into the specific regions of open chromatin that were made accessible by LA treatment, we proceeded with ATAC sequencing on LA treated MCF-10A cells. We examined the genomic locations of ATAC-seq peaks, representing open chromatin sites, and discovered 1704 open chromatin sites. Open chromatin regions were overrepresented within 1 kb of transcription start sites (TSSs) by 40-fold relative to the whole genome **(Fig. 2C**). Further, KEGG pathway analysis revealed 326 open chromatin regions with a log fold change >= 1.5 and FDR < 0.05 compared to vehicle treated cells. Among the top pathways that were upregulated significantly upon LA treatment are MAPK signaling pathway, PI3K-AKT signaling pathway, and the cAMP adenylate cyclase pathway. Additionally, motif analysis conducted using ‘HOMER’(*36*) showed that chromatin regions made accessible/inaccessible by LA treatment have binding motifs for a number of transcription factors (**Fig. 2E**). These data reveal that linoleic acid affects chromatin heterogeneity and increases/decreases the accessibility of specific regions that include transcription factor binding sites.

### Notch pathway genes are overexpressed in patients at high risk of ER-disease

Next, we sought to determine whether the genes, or sets of genes/pathways that we identified in our *in vitro* study were also differentially expressed *in vivo* in tissue of patients at risk for ER-and ER+ breast cancer. We took advantage of RNA from the contralateral unaffected breast (CUB) of breast cancer cases utilized in our previous studies, which revealed the association of LiMe genes in the CUBs of women with unilateral ER-breast cancer (*9, 10*). We combined the data from the RNA and ATAC sequencing experiments and collated a list of 44 genes of interest and 3 housekeeping genes. The list consists of the genes from the *HEDGEHOG, NOTCH, WNT, EMT, PPARγ* and adenylate cyclase pathways (**supplementary file S1**). TaqMan low density arrays were utilized to measure expression of these genes in CUBs of ER- and ER+ cases compared with the reduction controls. The study population included 84 women, with participants comprised of 28 matched triplets of women with ER-positive breast cancer, ER-negative breast cancer, and reduction mammoplasty controls. The three groups were matched by age, race and menopausal status as shown in **Fig. S2A**. As noted in our original publication, ANOVA revealed a significant difference in BMI across the three groups with BMI in the reduction mammoplasty control group (30.0 ± 5.8) notably higher than in ER-negative cases (25.3 ± 6.3, p=0.015), but not significantly higher than in the ER-positive group (26.7 ± 5.5, p= 0.136) (*10*). There was no significant difference in HER2 status between ER-positive and ER-negative cases. The majority of the selected genes had higher expression in high risk CUB specimens than the controls, irrespective of the ER status of the index tumor (**Fig. S2B**). The comparison between the ER- and ER+ CUBs revealed that in the ER-CUBS there is increased expression of genes that function in the Notch pathway: *NOTCH1* (1.7-fold, p=0.002, BH_adjP=0.07), *NOTCH4* (1.7-fold, p=0.04, BH_adjP=0.3), *DLL4* (2.5-fold, p=0.7, BH_adjP=0.8) and *HEY 1* (1.5-fold, p=0.05, BH_adjP=0.3), in addition to the *SMO* gene (1.47-fold, p= 0.05, BH_adjP=0.3), which is a key component of the hedgehog signaling pathway (**Fig. 3**). Altogether, these data reveal upregulation in *NOTCH* signaling in benign breast tissue samples from women at risk for ER-disease, suggesting that dysregulation of these pathways may play a role in the early stages of ER-cancer development.

**Fig. 3.**
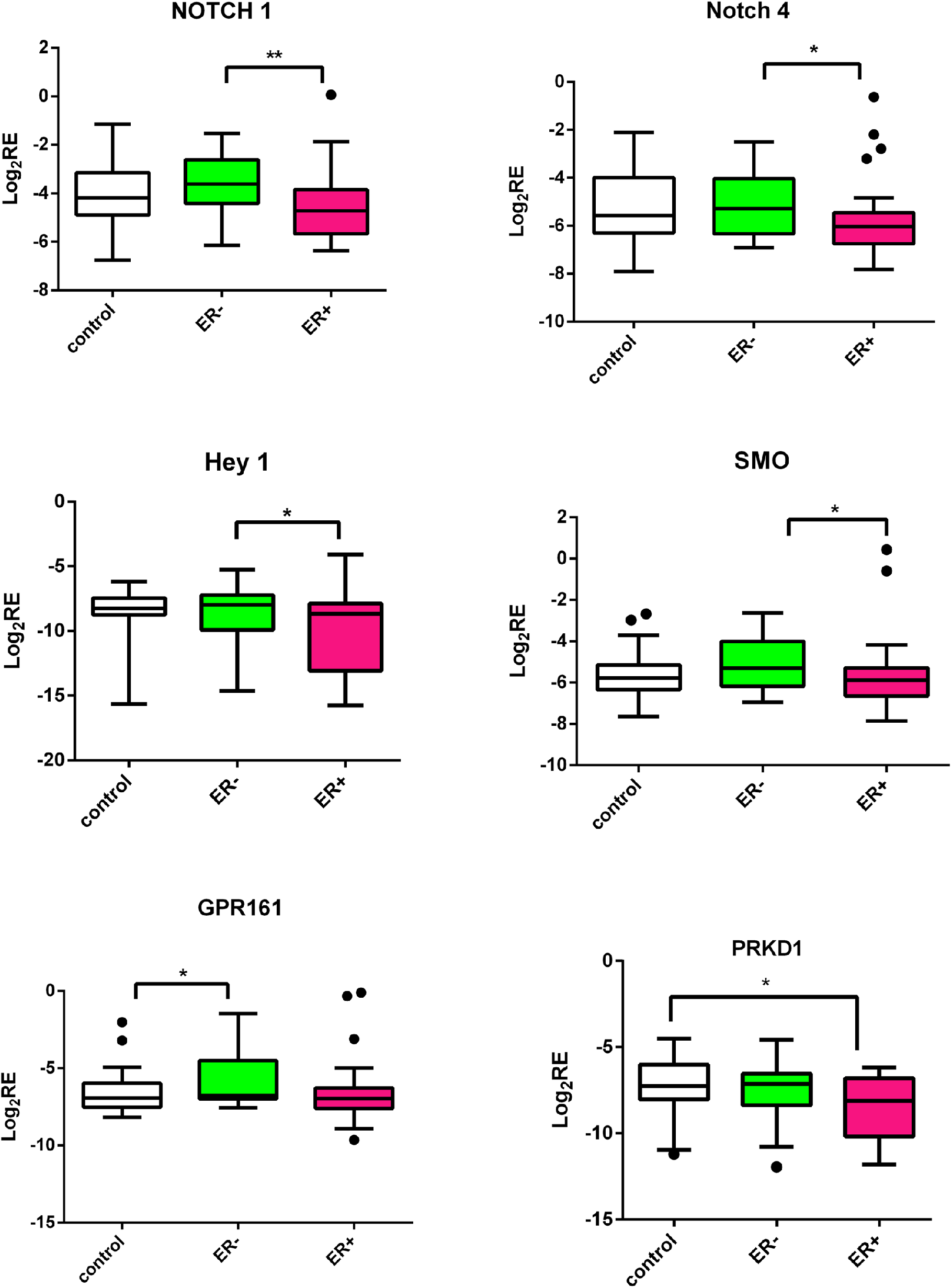
Notch pathway is overexpressed in CUB samples of patients at high risk of ER-disease. Expression of genes from various pathways in matching CUBs from ER negative, ER positive patients and controls. The log2-transformed relative (log2RE) amounts of mRNA expression normalized to the housekeeping gene and expressed as log_2_2^−(CtX−CtGAPDH^) = −(CtX − Ct GAPDH) where Ct is threshold cycle and X is gene of interest. IGF2 and GPR161 were significantly higher in ER negative versus normal whereas ER positive showed significant increase in PRKD1 versus normal. Genes from the Notch pathway were significantly higher in ER negative CUBs in comparison to ER positive patients. Mann-Whitney test was used to test the pairwise differences between the samples (ER+, ER-, Control) * p < 0.05; **p < 0.01.

### LA increases the expression of Notch pathway genes and specific genes involved in fatty acid oxidation *in vitro*

The increased expression of Notch pathway genes we discovered in the ER-CUBs, along with the similar findings in MCF10A cells exposed to octanoate (described above), led us to test the hypothesis that long chain fatty acids have similar effects on gene expression. We therefore investigated whether an increased LA environment influences the expression of Notch pathway genes and specific genes involved with fatty acid oxidation *in vitro*. We treated MCF-10A cells and mammary organoids from reduction mammoplasty patient samples with LA for 24 hours and then quantified changes in gene expression using RT-qPCR. To begin with, we assayed the genes involved in the activation of fatty acid oxidation. Upon entering cells, free fatty acids are converted into a fatty acyl-CoA molecules by the enzymes of the acyl-CoA synthetase (ACS) family (*37*). Notably, acyl-CoA synthetase long chain (*ACSL3*) is one of the LiME genes found to be upregulated in high risk ER-CUBs samples. Generation of acetyl-CoA occurs through a cyclical series of reactions in which a fatty acid is shortened by two carbons per cycle, eventually generating acetyl co-A. Acetyl co-A is a substrate for ketogenesis, which is initiated by the mitochondrial enzyme 3-hydroxy-3-methylglutaryl-CoA synthase 2 (*HMGCS2*), another of the previously identified LiMe genes. The mechanism for LCFAs oxidation is slightly more complex than for MCFAs, as this is regulated primarily via the enzyme carnitine palmitoyltransferase 1 (*CPT1*), the rate limiting enzyme of FAO which enables transport into the mitochondria. As shown in **Fig. 4A**, the expression of *HMGCS2, ACSL3*, and *CPT1B* were increased by LA exposure in MCF-10A cells and mammary organoids. Additionally, we observed a significant increase in *DLL4* expression followed by *HEY1, HEY2* and *NOTCH1* in the lipid treated mammary cells (**Fig. 4A**). We revisited the ATAC sequencing data to examine the effect of LA on chromatin architecture near key genes in the *DLL4*/*NOTCH* signaling pathway, and observed increased accessibility around the transcription start sites of *DLL4, NOTCH1* and *HEY1* showing significant lowered chromatin density with p-values of 1.62e-17, 0.017 and 0.03 respectively (**Fig. 4B & Fig. C**).

**Fig. 4.**
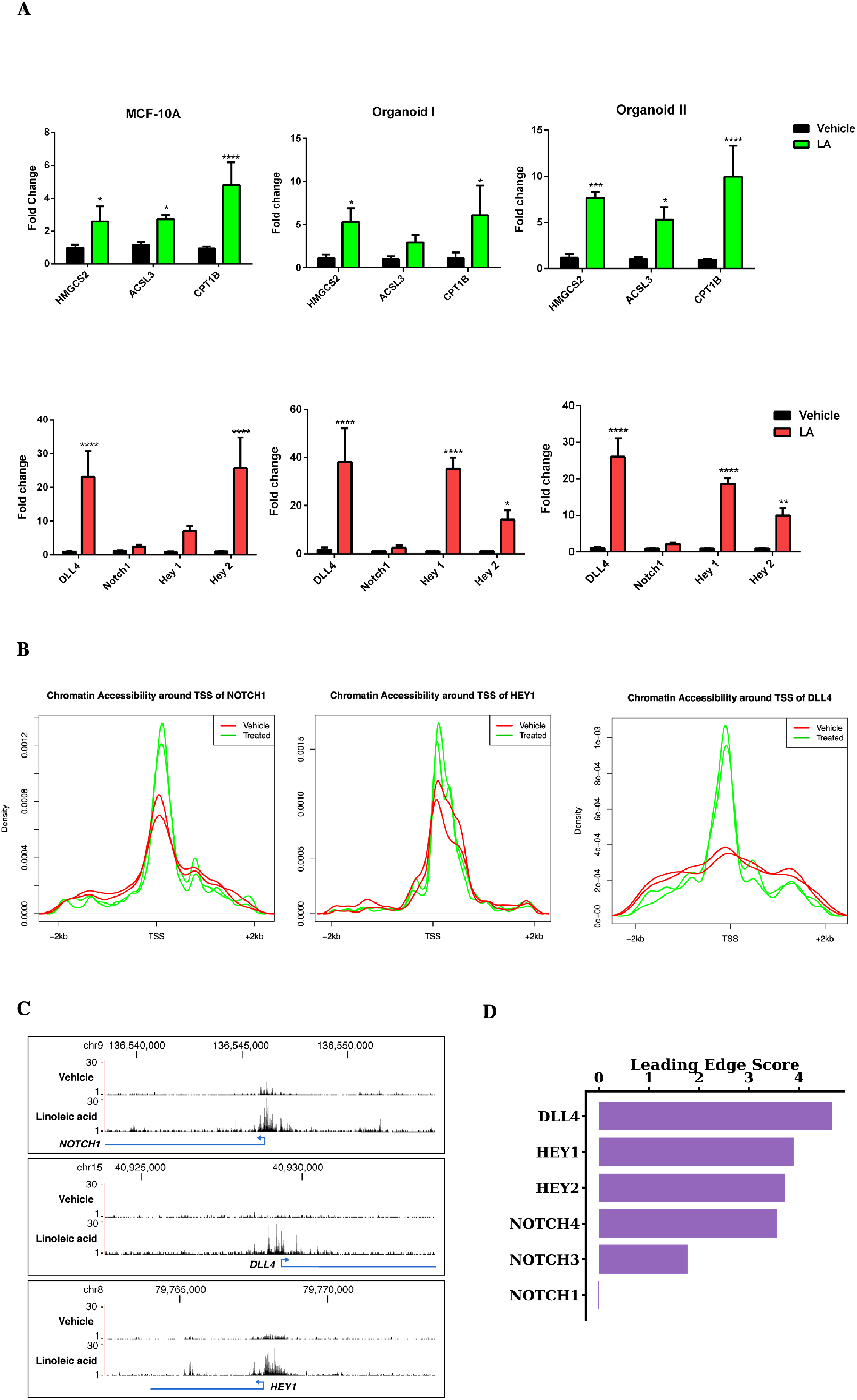
Increased DLL4/Notch signaling is associated with the stimulated fatty acid oxidation. (**A**) qPCR data showing increase in lipid metabolism genes (green) and Notch pathway genes (red) after 24-hour linoleate treatment in MCF-10A and mammary organoids. Statistical significance was determined by the unpaired t-test with Welch’s correction. (**B**) Chromatin accessibility in the lipid treated cells around the transcription start site (TSS) of NOTCH1, HEY1 and DLL4 (FDR<0.001). (**C**) Gene tracks and increase in peaks for the Notch genes in LA treated cells with the exact location on the chromosome. (**D**) Leading edge scores for genes of interest associated with the NOTCH signaling pathway as determined by GSEA leading edge analysis. DLL4, HEY1, HEY2, NOTCH3, and NOTCH4 were identified as core enrichment genes in the NOTCH pathway.

### Fatty acids drive flux through metabolic reactions resulting in increased histone methylation

While most of the experiments reported by McDonnell et al. were performed in AML 12 liver cells, these investigators also demonstrated increased H3K9 acetylation in octanoate-exposed MCF7 and MDAMB-231 breast cancer cells (*24*). Therefore, we sought to determine if these same experimental conditions would lead to H3K9 acetylation in a non-malignant MCF-10A cells. We exposed MCF-10A non-transformed ER - breast epithelial cell line to 5mM octanoate (C8) for 24 hours in medium containing both glucose (1.441 g/L) and glutamine (0.292 g/L). Western blot analysis demonstrated that octanoate exposure of MCF-10As resulted in increased acetylation at both H3K9 and H3K14 (**Fig. 5A**). To demonstrate that this was a fatty acid-specific effect, we treated the cells with 1,4-Cyclohexanedimethanol (1,4-CHDM), an alcohol with the same formula as octanoate; no acetylation was observed consequent to the alcohol exposure (**Fig. S3A**). To validate the specificity of the antibody against the acetylated histone lysines, we treated MCF-10A cells with sodium butyrate, a histone deacetylase (HDAC) inhibitor. Sodium butyrate treatment increased the acetylation of H3K9 and H3K14 as shown in **Fig. S3B**.

**Fig. 5.**
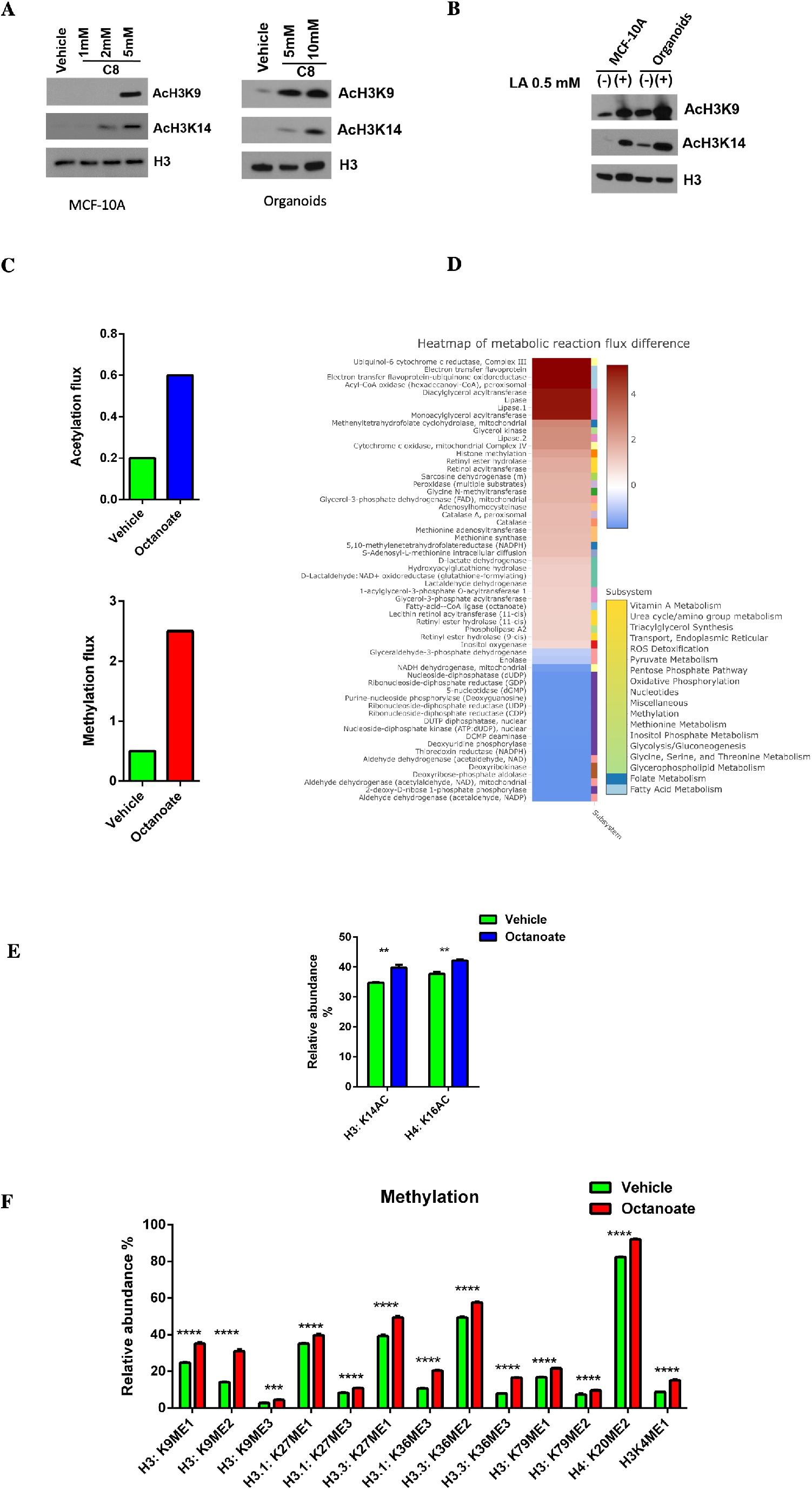
Fatty acids drive histone modifications and metabolic flux. Western blot of histone acetylation at H3 lysine K9 and K14 in MCF-10A cells and organoids treated with (A) octanoate and (B) linoleic acid. **(C)** The effect of octanoate treatment on histone acetylation and methylation flux in MCF-10A cells predicted using genome-scale metabolic modeling. (D) Heatmap of reaction flux differences predicted by metabolic modeling to be differentially active (p-value < 0.01) between control and treatment. The corresponding pathways (subsystem) that each reaction belongs to is listed in the legend. Proteomic acetylation (E) and methylation (F) profiling measured by mass spectrometry of MCF-10A cells treated in triplicate with 5mM octanoate for 24 hours in a complete media compared to vehicle. Two-way ANOVA was performed to determine the statistical significance and corrected for multiple comparisons using Sidak test.

**Fig. 6.**
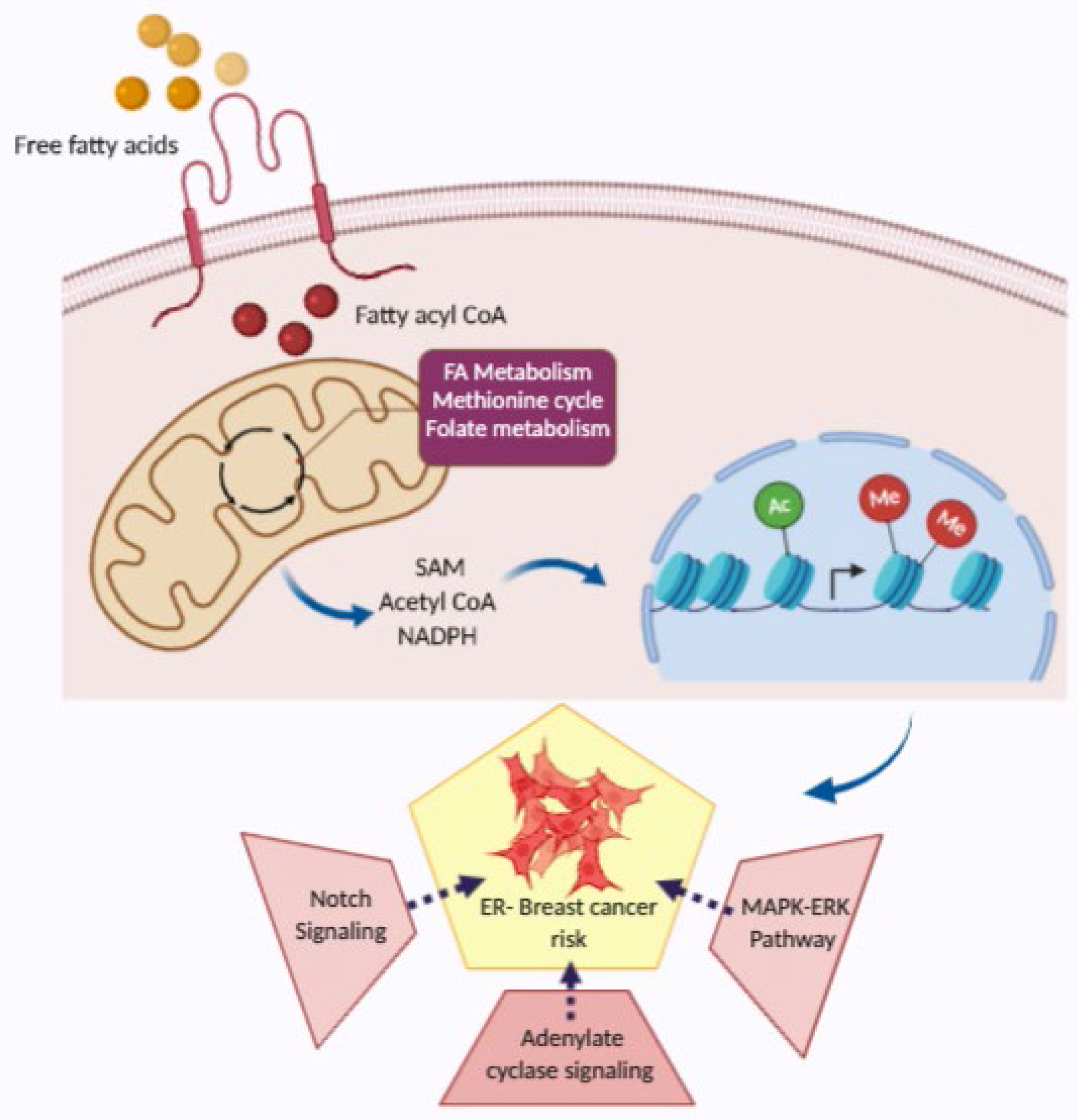
Proposed model illustrating the orchestration of lipid induced molecular changes. *Sensors*: Senses the fatty acid rich environment and perturb cellular metabolism providing the essential substrate for histone modifications and thereby turning on the *Mediators-*histone PTMs, which consequently activates the *Effectors-*Notch, adenylate cyclase and MAPK-ERK the key protein signaling associated with ER-breast cancer.

To exhaustively explore the impact of octanoate treatment on metabolic pathways, we used flux balance analysis (FBA) (*38*). FBA makes use of genome-scale metabolic network models that contain all known metabolic reactions in a cell or tissue based on evidence from the published literature (*39*). Genome-scale metabolic models have been widely used to predict the metabolic behavior of various mammalian cell types (*40-44*). Here we used the Recon1 human network model that maps the relationship between 3744 reactions, 2766 metabolites, 1496 metabolic genes, and 2004 metabolic enzymes (*45*). This model was augmented with biochemical reactions corresponding to histone acetylation and methylation (*40, 46*), allowing us to predict the consequences of octanoate-induced metabolic changes on histone modifications by tracking the flux through the substrates for the histone modifications. These models were previously used to predict bulk histone acetylation levels in various cell lines based on the nuclear flux of acetyl-coA directed towards histone acetylation (*46*). Similarly, bulk histone methylation levels can be predicted based on the nuclear flux of S-adenosyl-L-methionine (SAM) (*40*).The model predicted octanoate treatment would result in increased histone methylation levels, with a more modest increase in histone acetylation levels (**Fig. 5C**). As a comparison, we repeated this analysis with immortalized hepatocyte cells used by McDonnel et al; they found a significant increase in histone acetylation after octanoate treatment (*24*). We calculated metabolic flux in these hepatocytes using the transcriptomics data from McDonnel *et al* and found a much larger increase in histone acetylation after octanoate treatment (**Fig. S3C**). These results suggest that the impact of metabolic alterations on histone acetylation is cell-type specific, as observed in prior studies (*47, 48*). Overall, out of the 3759 reactions in the model, we identified 38 that showed significant increased activity after octanoate treatment (p-value < 0.01; **Fig. S3C**). As expected, reactions involved in lipid and fatty acid metabolism, specifically triacyl glycerol synthesis and glycerophospholipid metabolism were upregulated. Interestingly, among the up-regulated reactions were several reactions related to the one-carbon metabolic pathway, which links folate, SAM, methionine, glycine and serine metabolism (**Fig. 5D**). The reactions methionine adenosyltransferase, methionine synthase, adenosyl homocysteinase, 5,10-methylene-tetrahydrofolatereductase, glycine N-methyltransferase, and formyltetrahydrofolate dehydrogenase were all predicted to have increased activity after treatment (p-value < 0.01). Increased activity of the one-carbon pathway is associated with increased H3K4 trimethylation in stem cells and cancer cell lines (*40, 49*). These reactions likely support increased histone methylation by providing one carbon units.

### Lipid exposure eventuates in histone methylation

In order to profile the specific histone marks significantly changed by the octanoate treatment we performed liquid chromatography/mass spectrometry on tryptic peptides isolated from the nuclei of treated and control MCF10A cells. Increased methylation was observed in various histone proteins including H3K9me1/2/3, H3.1K27me2/3, H3.3K36me2/3, H3K79me1/2 and H3K4 (**Fig. 5F**) together with increased acetylation of H3K14 and H4K16 (Fig. 5E). Notably, the GSEA analysis showed a significant correlation of H3K27 methylation (NES = 2.47, FDR q-value =0.05) and H3K4 methylation (NES= 1.24, FDR q-value = 0.1) with octanoate treatment (**Fig. S3 D-E**) suggesting this lipid rich environment eventuates in histone methylation in mammary epithelial cells.

## Discussion

The known determinants of risk for ER-negative breast cancer are genetic (either specific racial inheritance, germline mutations in genes such as BRCA1) or systemic/behavioral factors (premenopausal obesity (*50*), absence of a breastfeeding (*51*)). In contrast, few if any local factors in the breast environment serve to identify women at risk for ER negative tumors. Local in-breast factors are of great interest however, since they may be more specifically targetable for breast cancer prevention than systemic factors. Of note, the two strongest risk factors for breast cancer overall (other than high penetrance germline mutations) are local: atypical proliferative lesions, and (*52*) extremely dense breast tissue (*53*). This reasoning motivated us to investigate the local breast biology that may promote the development of ER negative rather than ER positive breast cancer, using the contralateral unaffected breast (CUB) of women undergoing surgery for a unilateral primary breast cancer as a model for ER-specific breast cancer risk (*7, 54*). In our initial study, we identified a highly correlated lipid metabolism (LiMe) gene signature, which was enriched in the CUBs of women with ER-breast cancer.

To explain the biologic basis for this association, we developed an *in vitro* model wherein we exposed either MCF10A, an ER negative, non-tumorigenic epithelial cell line, or breast organoids derived from reduction mammoplasty samples to an extracellular milieu rich in medium or long chain fatty acids. This model system has now enabled us to demonstrate that the exposure of breast epithelial cells to these fatty acids results in a dynamic and profound change in gene expression, accompanied by changes in chromatin packing density, chromatin accessibility and histone PTMs. The histone modifications, in turn, are the result of both the lipid-engendered increased expression of the requisite enzymes and the increased production of their substrates. Our metabolic flux analysis revealed the upregulation of several reactions related to the one-carbon metabolic pathway, which links folate, SAM, methionine, glycine and serine metabolism. This insight was not evident upon analysis of differential gene expression, which is not surprising as gene expression changes often do not reflect the flux of metabolic reactions (*40*).

Our proteomics data reveal increased methylation at H3K27me2/3, H3K36me3 and H3K9me2/3 in cells treated with octanoate; GSEA analysis showed that genes with ontologies related to histone methylation at H3K27 and H3K4 exhibit changes in expression in the lipid-treated cells. Methylation of H3K27 is carried out by *EZH2*, which showed a 1.65-fold increase in expression (p=0.001) following exposure to octanoate. *EZH2* expression is sensitive to the level of oxidative phosphorylation. Our metabolic flux data demonstrated increased oxidative phosphorylation following exposure to octanoate; specifically flux through the electron transport chain (ETC). Inhibition of oxidative phosphorylation via complex I of the ETC by the biguanide phenformin markedly reduces EZH2 and SUZ12 protein expression (*55*). This suggests that increased H3K27 methylation may be a consequence of increased flux through the ETC increasing EZH2 expression in concert with increased production of its substrate SAM. Several studies have revealed a significant association of *EZH2* overexpression with ER negative breast cancer (*56*) or ER negative luminal progenitor cell expansion (*57*). EZH2 is the enzymatic subunit of Polycomb Repressive Complex 2, which catalyzes the trimethylation of H3K27. However, EZH2’s actions are not be limited to its methyltransferase activity. EZH2 has been shown to bind to the *NOTCH 1* promoter resulting in increased *NOTCH1* transcription, stem cell expansion and accelerated tumor initiation (*58*). The effect of *NOTCH1* expression on mammary cell-lineage fate determination was recognized shortly after the identification of the mammary stem cell (*59*). Mammary stem cell differentiation is a hierarchical organization, and lineage tracing experiments have determined that *NOTCH1* expression exclusively generates ER-luminal cells (*60*). A subsequent study by these investigators revealed that during mammary embryogenesis Notch signaling prevents the generation of basal precursors, and cells expressing active *NOTCH1* exclusively give rise to the ER-(Sca1-/CD133-) lineage at any developmental stage from mouse embryonic day 13.5 to postpartum day 3 (*61*). Even more interesting given our focus on the origins of ER negative breast cancer was their observation that pubertal cells retain plasticity. Ectopic activation of Notch1 in basal cells at puberty was able to completely switch their identity to ER negative luminal cells.

Additional clues regarding the association of our experimental findings with ER negative breast cancer comes from GWAS data. A study that included 21,468 ER-negative cases and 100,594 controls identified independent associations of ten single nucleotide polymorphisms (SNPs) with the development of ER-breast cancer (*62*). Pathway analysis was performed by mapping each SNP to the nearest gene. This identified several pathways implicated in susceptibility to ER-negative, but not ER+ breast cancer. Included among these was the adenylate cyclase (AC) activating pathway. One of the significantly altered biologic processes that we identified by RNA sequencing of the octanoic acid treated cells is adenylate cyclase-activating adrenergic receptor signaling. Adenylate cyclase signals via cyclic AMP. Regions of chromatin with increased accessibility are associated with increased gene expression; our ATAC-Seq results show that linoleic acid exposure significantly increased accessibility to genes in the cAMP signaling pathway. In their discussion of ER-GWAS results, Milne et al. suggest that stimulation of the beta 2 adrenergic-adenylate cyclase-cAMP-β-arrestin–Src–ERK pathway may play a role in the genesis of ER-breast cancer. MetaCore analysis of our RNA-sequencing data reveals similar pathway activation, however, it is the beta1 adrenergic receptor that demonstrates increased expression in the octanoate treated cells. In addition, our ATAC-seq data showed increased RAP1 signaling pathway accessibility. Adenylate cyclase signaling also functions via Epac-Rap1-B-raf-MEK-ERK, with this signaling shown to be responsible for sustained *ERK* activation that occurs at a later time points (10-30 minutes) after cAMP activation (*63*). The MAPK (ERK) pathway can be stimulated by means other than adrenergic receptor ligand binding. Activation of this pathway by overexpression of *EGFR*+*EGF, c-erbB-2, RAF1* or *MEK* in MCF7 cells leads to estrogen-independent growth and down-regulation of ERα expression (*64*). These results suggest that hyperactivation of the MAPK(ERK) pathway plays a role in the generation of the ER-phenotype in breast cancer. We observed *MAPK* activation in our analysis of differentially expressed genes, i.e., “positive regulation of the *MAPK* cascade,” and in the analysis of regions of chromatin with significantly increased chromatin.

Using stratified LD score regression, a statistical method for identifying functional enrichment from GWAS summary statistics, SNPs associated with the H3K4me3 histone mark were determined to be contributing to the heritability of ER-negative breast cancer, (2.4-fold, P = 0.0005) (*62*). Increased activity of the one-carbon pathway is associated with increased H3K4 trimethylation in stem cells and cancer cell lines (*40, 49*). Restriction of methionine with consequent modulation of SAM and S-Adenosyl-L-homocysteine (SAH) levels affects methylation at H3K4me3, H3K27me3 and H3K9me3, with H3K4me3 exhibiting the largest changes (45). Interestingly, this restriction leads to loss of H3K4me3 at the promoters of colorectal cancer (CRC)-associated genes, with resulting decreased expression (p = 0.02, Fisher’s exact test). A computational model developed to identify the direct influences on methionine concentrations in humans suggests that dietary intake explains about 30% of the variation in methionine concentration, and fats (arachidic acid in this model) are among the foods contributing to higher methionine levels (*49*).

One-carbon metabolism has multiple other functions in addition to producing SAM; one of which is to maintain redox homeostasis by producing NADPH. One of the earliest steps in breast tumorigenesis is the filling of the duct/acinar lumen with malignant cells. The viability of ECM-detached cells is dependent on combating the generation of reactive oxygen species (ROS) (*65*). For example, shuttling flux through the pentose phosphate pathway (PPP) promotes NADPH production and consequent reduction of ECM detachment-induced reactive oxygen species. It appears, however, that the process that produces the reducing equivalent is immaterial as cells can also utilize NADPH-regenerating enzymes in the folate pathway as in metastasizing melanoma (*66*). Therefore, the one-carbon metabolism initiated by FAs may facilitate early tumorigenesis and the survival of matrix detached cells by the production of NADPH.

In conclusion, we have demonstrated in the present study that exposure of breast epithelial cells *in vitro* to fatty acids results in epigenetic effects that produce dynamic and profound changes in the expression of genes that have been associated with the development of ER-breast cancer. Next steps include demonstrating that these same changes are observed *in vivo*. As mentioned in the introduction, polyunsaturated fatty acids are present in normal breast tissue. Although we measured lipid species in the serum of the donors of the CUB specimens, fatty acids can also be mobilized from adjacent adipose tissue; adipocytes have been shown to be a reservoir of lipids for breast cancer stem cells (*67*). We hypothesize that the expression of genes associated with the development of ER-breast cancer is consequent to lipid stimulation of one-carbon metabolism with resultant changes in histone methylation. Important roles for glycolysis, glutaminolysis, lipogenesis and mitochondrial activity have been demonstrated in oncogenesis; the one-carbon pathway has comparatively received less attention and the insights we provide here generate new questions regarding lipid metabolism and ER negative breast cancer, to be pursued in future investigations.

## Materials and Methods

### Cell culture

MCF10A cell line was obtained from American Type Culture Collection (ATCC) and cultured in mammary epithelial cell growth basal medium with single quots supplements and growth factors (#Lonza CC-4136). Cells were treated with the medium-chain fatty acids (Sigma) sodium octanoate (C8) dissolved in PBS, and long-chain fatty acids (Sigma) Linoleic acid (C18) complexed with fatty acid free BSA (Roche 10775835001). PBS and BSA were used as the vehicle control in experiments containing C8 and C18 respectively. Cells were counted using an Invitrogen Countess automated cell counter using Trypan blue exclusion method and seeded at the indicated densities. All experiments were done in complete MEBM media with fatty acids or vehicle.

### CUB Samples

Patients diagnosed with unilateral breast cancer and undergoing contralateral prophylactic mastectomy at Prentice Women’s Hospital of Northwestern Medicine were recruited under an approved protocol (NU11B04), with exclusions for neoadjuvant treatment, prior endocrine therapy or pregnancy/lactation during the prior 2 years. A group of reduction mammoplasty (RM) patients were also recruited as standard risk controls. The fresh tissues were frozen and stored in liquid nitrogen. Tissue samples from 56 bilateral mastectomy cases (28 ER+ and 28 ER–) and 28 healthy RM controls were used in this study. The ER+ cases, ER– cases and controls were matched by age, race, and menopausal status.

### Mammary Organoids Preparation

Tissues were collected from the non-obese, premenopausal women coming for the reduction mammoplasty. Transfer the breast tissue to be processed into a sterile petri dish. Chop big breast tissue mass into small pieces. Transfer the minced tissue to a sterile 50ml tube and add 30ml of Kaighn’s Modification media (Gibco #21127022) containing collagenase from Clostridium histolyticum (Sigma Aldrich, catalog no. C0130), final collagenase concentration is 1 mg/mL. Media containing collagenase is filtered using 0.22 μm filter. The falcon is sealed with parafilm and tissue is gently dissociated on a shaker at 100 rpm and 37°C, overnight (16 hours). Following day, organoids are collected by the centrifugation of the suspension at 800 rpm for 5 min. Discard the supernatant and wash the organoid pellet two-three times with PBS. Organoids with a size between 40-100uM are collected and resuspended in the fresh media (3mL) and added to a 6 well plate (Ultra-Low Attachment Surface plate, Corning # CLS3471). Organoids are allowed to stabilize for 24 hours before using it for the experiments.

### Fatty acid preparation

Sodium octanoate (C8) was dissolved in PBS. To bind linoleic acid (Sigma # L8134) to BSA, they were initially dissolved in water to yield a 50 mM final concentration. Dissolve 0.12g of BSA in 1.2 ml of water resulting a 10% BSA solution. Combine 0.2 ml aliquot of the Na linoleate solution to the 10% BSA solution. After 15 min of slow stirring at 37°C, 0.6 ml of water was added to bring the final concentration of Na linoleate to 5 mMol/L (Pappas et al, 2001).

### Lipid analysis

LC-MS grade methanol, dichloromethane, and ammonium acetate were purchased from Fisher Scientific (Pittsburgh, PA) and HPLC grade 1-propanol was purchased from Sigma-Aldrich (Saint Louis, MO). Milli-Q water was obtained from an in-house Ultrapure Water System by EMD Millipore (Billerica, MA). The Lipidyzer isotope labeled internal standards mixture consisting of 54 isotopes from 13 lipid classes was purchased from Sciex (Framingham, MA).

### Sample Preparation

Frozen plasma samples were thawed at room temperature (25 °C) for 30 min, vortexed; 25 uL of plasma was transferred to a borosilicate glass culture tube (16 x 100 mm). Next, 0.475 mL of water, 1.45 mL of 1:0.45 methanol:dichloromethane, and 25 uL of the isotope labeled internal standards mixture were added to the tube. The mixture was vortexed for 5 sec and incubated at room temperature for 30 min. Next, another 0.5 mL of water and 0.45 mL of dichloromethane were added to the tube, followed by gentle vortexing for 5 sec, and centrifugation at 2500 g at 15 °C for 10 min. The bottom organic layer was transferred to a new tube and 0.9 mL of dichloromethane was added to the original tube for a second extraction. The combined extracts were concentrated under nitrogen and reconstituted in 0.25 mL of the mobile phase (10 mM ammonium acetate in 50:50 methanol:dichloromethane).

### Mass Spectrometry

Quantitative lipidomics was performed with the Sciex Lipidyzer platform consisting of Shimadzu Nexera X2 LC-30AD pumps, a Shimadzu Nexera X2 SIL-30AC autosampler, and a Sciex QTRAP® 5500 mass spectrometer equipped with SelexION® for differential mobility spectrometry (DMS). 1-propanol was used as the chemical modifier for the DMS. Samples were introduced to the mass spectrometer by flow injection analysis at 8 uL/min. Each sample was injected twice, once with the DMS on (PC/PE/LPC/LPE/SM), and once with the DMS off (CE/CER/DAG/DCER/FFA/HCER/LCER/TAG). The lipid molecular species were measured using multiple reaction monitoring (MRM) and positive/negative polarity switching. Positive ion mode detected lipid classes SM/DAG/CE/CER/DCER/HCER/DCER/TAG and negative ion mode detected lipid classes LPE/LPC/PC/PE/FFA. A total of 1070 lipids and fatty acids were targeted in the analysis.

### Data Processing

Data was acquired and processed using Analyst 1.6.3 and Lipidomics Workflow Manager 1.0.5.0. For statistical analysis, we evaluated the lipid species enrichments in the ER+, ER-, and control groups. The different groups were compared in pair-wise and the log-fold changes of lipid enrichment were derived, along with the effect sizes and p-values inferred from the regression models using the lipid measurement as an input variable and group information as the output variable.

### Library preparation and RNA Sequencing

RNA was isolated with Qiagen RNeasy Plus Mini Kit (# 74134) as per the manufacturer’s protocol. The concentration and quality of total RNA in samples were assessed using Agilent 2100 Bioanalyzer. RNA Integrity Number (RIN) of the vehicle and octanoate sample was 9.9 and 9.8 respectively. Sequencing libraries were prepared from a total of 100ng of RNA using KAPA RNA HyperPrep Kit. Single-Indexed adapters were obtained from KAPA (Catalog# KK8701). Library quality was assessed using the KAPA Library Assay kit. Each indexed library was quantified and its quality accessed by Qubit and Agilent Bioanalyzer, and 6 libraries were pooled in equal molarity. 5μL of 4nM pooled libraries were denatured, neutralized and a final concentration of 1.5 pM of pooled libraries was loaded to Illumina NextSeq 500 for 75b single-read sequencing. Approximately 80M filtered reads per library was generated. A Phred quality score (Q score) was used to measure the quality of the sequencing. More than 88% of the sequencing reads reached Q30 (99.9% base call accuracy). Single-end FASTQ reads from RNA-seq measurements were aligned and mapped to hg38 ENSEMBL genome using STAR alignment (*68*).

### Gene Ontology Analysis of Differentially Expressed Genes

Transcriptions per million (TPM) from mapped reads were estimated using RSEM from the STAR aligned reads (*69*). The *DESeq2* R package (*70*) was employed to determine differentially expressed genes for the octanoate treatment group compared to the vehicle-treated controls with FDR cutoff = 0.01 and |log_2_ *FC*| ≥ 2 to identify a reasonable number of differentially expressed genes, on the order of several thousands of genes total, for subsequent analysis. Gene ontology pathway analysis for biological processes was performed on each set of differentially expressed genes using *Metascape* (*71*).

### GSEA Analysis

Raw counts were first estimated using *HTSeq* from STAR aligned reads *(72)*. Next, replicates for control cells and treated cells were merged and normalized using modules from the GenePattern software package (*73*). Gene set enrichment analysis (GSEA) (*74, 75*) was performed on these DESeq-normalized reads using annotations from online databases, including KEGG, Hallmark, Reactome, BioCarta, and Canonical Pathways. The normalized enrichment score (NES) of these top 20 pathways associated with the control and the octanoate-treated condition are shown with nominal p-value = 0.0. *Metascape* was employed to perform network analysis on these top 20 pathways associated with each treatment condition.

### ATAC Seq Library preparation and sequencing

1×10^6^ cells were pelleted and lysed in ATAC-resuspension buffer as described (*76*). Extracted nuclei was processed for TN-5 mediated tagmentation using the Illumina Tagment DNA Enzyme and buffer kit (Nextera Illumina # 20034210) : Transposon reaction mix as 2X TD Buffer-25 µl, Tn5 Transposase – 2.5µl, 1X PBS containing nuclei-16.5µl, 10% Tween-20-0.5µl (Sigma # P9416), 1% Digitonin-0.5µl (Promega # G9441) and water at 37°C, 1000rpm for 30mins. Tagmented DNA was isolated by Nucleospin PCR clean-up (Takara Bio USA, Inc # 740609.250). Libraries were amplified for 8 cycles and purified using AMPure XP (Agencourt # A63880). Fragment sizes were determined using 106 LabChip GXII Touch HT (PerkinElmer), and 2×50 paired-end sequencing performed on NovaSeq S1 6000 flow cell (Illumina) flow to yield 100M reads per sample.

### ATAC-seq data sequencing and peak calling

Illumina adapter sequences and low-quality base calls were trimmed off the paired end reads with Trim Galore v0.4.3. Sequence reads were aligned to human reference genome hg38 using bowtie2 with default settings. Duplicate reads were discarded with Picard. Reads mapped to mitochondrial DNA together with low mapping quality reads were excluded from further analysis. MACS2 was used to identify the peak regions with options -f BAMPE -g hs –keep-dup all -B -q 0.01. Peaks for samples in the same condition were merged using the function ‘merge’ of bedtools and peaks for samples in different conditions were intersected using the function of ‘intersect’ of bedtools.

### Differential chromatin accessibility analysis

The number of cutting sites of each samples were counted using the script dnase_cut_counter.py of pyDNase. The raw count matrix was normalized by CPM. R package edgeR was used to conduct the differential accessibility analysis for all 66,853 common peaks. Significant different accessible chromatin regions under different conditions were defined as the threshold 0.05 for FDR. With the cutoff 1 for the absolute value of fold change, comparing treatment group with vehicle control group, we got 1,704 significant increased peaks and 3,340 significant decreased peaks.

### Motif analysis

Motif analysis were conducted for significant changed chromatin regions using ‘findMotifsGenome.pl’ script of HOMER with default settings. The principal component analysis was conducted to detect the important motifs using the relative enrichment of motifs. Biplot was used to visualize the principal component analysis results.

### Genomic distribution of open chromatin regions

We calculated the overall genomic distribution of open chromatin regions, comparing the treatment to the vehicle, based on the methods as described (*77*). We used the hg38 refseq genes annotation from UCSC genome browser to define the genomic features. All TSSs were considered in the analysis if a gene had multiple TSSs. The formula for reported enrichment is (a/b)/(c/d). a is the number of peaks overlapping a given genomic feature, b is the number of total peaks, c is the number of regions corresponding to the feature, and d is the estimated number of discrete regions in the genome where the peaks and feature could overlap. Specifically, d is equal to (genome size)/ (mean peak size + mean feature size), following the implementation in the bedtools fisher.

### Pathway analysis for open chromatin regions

For the 326 open chromatin regions with logFC ≥ 1.5 and FDR < 0.05 comparing the treatment with the vehicle, we used R package ‘clusterProfile’ to conduct KEGG pathway analysis.

### Validation of candidate genes qRT-PCR

Treated cells and organoids were washed with PBS and RNA was isolated with Qiagen RNeasy plus mini Kit (# 74134) as per the manufacturer’s protocol. cDNA was synthesized using the SuperScript VILO cDNA synthesis kit (#11755250). Real-time qPCR was performed using Applied biosystem Quant studio 5 real time PCR System (Thermo Scientific). Expression data of the studied genes was normalized to RPLP1 to control the variability in expression levels and were analyzed using the 2^-ΔΔCT^ method described by Livak and Schmittgen (*78*). TaqMan gene expression assays were purchased from ThermoFisher Scientific and the list of the assays is provided in supplemental file **S1**.

### qRT-PCR based TaqMan low density array assays

Based on histological diagnosis atypical hyperplasia benign breast epithelium was identified and captured by laser capture microdissection (LCM). RNA was isolated with Qiagen RNeasy plus mini Kit (# 74134) as per the manufacturer’s protocol. RNA quality was checked for integrity using Bioanalyzer 2100 by Agilent. 100ng RNA was reverse transcribed using High Capacity RNA-to-cDNA Master Mix (#4388950) and preamplified for 14 cycles using TaqMan PreAmp Master Mix 2X((#4488593) and pooled assay mix for the genes in which we were interested. Pre-amplified cDNA were diluted to 1:20 with 1X TE buffer and mixed with Fast advanced master mix (# 4444965) Each sample was loaded in duplicate in 384-well microfluidic cards customized with 47 genes of interest including three housekeeping genes (GAPDH, RPLP0 and RPLP1). TaqMan assays with best coverage attribution were used for the TLDA study as recommended by the manufacturer. A list of the genes and the Assay ID for the primers obtained from ThermoFisher is provided in supplemental file S2. Real Time PCR reactions were carried out in Quant studio 7 Flex system for 40 cycles using comparative Ct (ΔΔCt) method. Results were analyzed using Expression suite software.

### Live cell PWS Imaging

Before treatment and imaging, MCF-10A cells were seeded in 6 wells black culture plate at least 35% confluency and allowed to adhere overnight before the treatment with 500µM LA (C18:2) and 5mM Octanoate. We based the concentration of LA used in the experiment on the range in human plasma: 0.2 to 5.0 mmol/L (*79*). For chromatin study experiments, live-cell PWS images were acquired at room temperature (22 °C) and in trace CO2 (open air) conditions. Imaging was performed using the commercial inverted microscope (Leica DMIRB) Hamamatsu Image-EM CCD camera C9100-13 coupled to a liquid crystal tunable filter (LCTF; CRi Woburn, MA) to acquire mono-chromatic spectrally resolved images that range from 500–700 nm at 1 nm intervals produced by a broad band illumination provided by an Xcite-120 LED Lamp (Excelitas, Waltham, MA) as previously described (33, 34). Briefly, PWS measures the spectral interference resulting from internal light scattering structures within the cell, which captures the mass density distribution. To obtain the interference signal directly related to refractive index fluctuations in the cell, we normalized measurements by the reflectance of the glass medium interface, i.e., to an independent reference measurement acquired in an area without cells. PWS measures a data cube (spatial coordinates of a location within a cell and the light interference spectrum recorded from this location). The data cube then allow to measure spectral SD (Σ), which is related to the spatial variations of refractive index within a given coherence volume. The coherence volume was defined by the spatial coherence in the transverse directions (∼200 nm) and the depth of field in the axial direction (∼1 mm). In turn, the spatial variations of refractive index depended on the local autocorrelation function (ACF) of the chromatin refractive index. Finite-difference time-domain simulations have shown that PWS is sensitive to ACF within the 20-to 200-nm range. According to the Gladstone-Dale equation, refractive index is a linear function of local molecular crowding. Therefore, S depends on the ACF of the medium’s macromolecular mass density. Small molecules and other mobile crowders within the nucleus are below the limit of sensitivity of PWS, and PWS is primarily sensitive to chromatin conformation above the level of the nucleosome. To convert S for a given location within a nucleus to mass fractal dimension D, we modeled ACF as a power law B_¥_(r)= ðrÞ ¼s2ϕrrmi_ _D_3, where ϕ is the variance of CVC (60). In general, S is a sigmoidal function of D. However, for fractal structures such as a chromatin packing domain where within physiological range 2 < D < 3, S can be approximated as a linear function of D by the relationship D ≈ D0 + aS, where D0 = 1.473 and is comparable to the minimal fractal dimension that an unconstrained polymer can attain and constant a ∼ 7.6. The measured change in chromatin packing scaling between treatment conditions was quantified by first averaging D within each cell’s nucleus and then averaging nuclei from over 100 cells per condition.

### Flux based analysis (FBA)

We calculated the relative activity of reactions in MCF-10A cells by interpreting gene expression data using the Recon1 human metabolic model augmented with histone modifications (*46, 80*). We then identified a metabolic flux state that is most consistent with gene expression data in control and octanoate treatment. This was achieved by maximizing the activity of reactions that are associated with up-regulated genes and minimizing flux through reactions that are down-regulated in a condition, while simultaneously satisfying the stoichiometric and thermodynamic constraints embedded in the model using linear optimization (*46, 80*). The glucose, fatty acid, and glutamine levels in the simulations were adjusted based on the growth media used for culturing the cells. All p-values were corrected for multiple comparisons.

### Statistical analysis

Prior to performing the analyses, the log2-transformed relative (log2RE) amounts of mRNA expression normalized to GAPDH and expressed as log_2_2^-(CtX- CtGAPDH)^ = -(CtX-CtGAPDH) where Ct is threshold cycle. Mann-Whitney test was performed to identify genes with pairwise differences between ER+ and ER- samples. The analyses were adjusted for multiple testing, 34 genes, using the Benjamini-Hochberg (BH) adjustment in order to control the false discovery rate at the two-sided 0.05 level. Boxplots were used to visualize differences in log2RE by group. The log2RE analyses were conducted using the R statistical environment [R] version 3.5.1.

## Supplementary Materials

**S1 List of primers used for qRT-PCR validation of candidate genes**. List of Assay ID (ThermoFisher) of primers utilized in qRT-PCR of candidate genes (Fig. 4A).

**S2 List of genes assayed by TaqMan low density array (TLDA) and their corresponding primers**. List of genes and the corresponding Assay ID (ThermoFisher) of primers used for TLDA assays.

## General

We would like to thank Professors Matthew D Hirschey and Neil Kelleher for advice regarding histone proteomics and, Jeannie Camarillo and the Northwestern Proteomics Core for conducting the histone proteomic analysis; The Northwest Metabolic Research Center (NW-MRC) at University of Washington for performing the lipidomics analysis; The Center for Medical Genomics at the Indiana University School of Medicine for RNA library preparation and RNA sequencing, and ATAC sequencing; The NU Seq Core facility for providing The Quant Studio 7 Flex system; Natalie Pulliam for consenting patients and collecting tissue, and our many lab colleagues for feedback.

## Author contributions

Conceptualization and project design: S.Y., S.E.C., and S.A.K.; experiments: S.Y ; Sequencing: X.X and H.G; RNA seq analysis : R.K.A.V., Z.Z., G.C and M.R.; ATAC Seq analysis: D.C.; PWS microscopy: D.V.D.; Metabolic flux analysis: C.H.C and S.C; Statistical Analysis: K.L.B ; writing (original draft): S.Y., S.E.C., and S.A.K.; writing (review and editing): V.B., S.C., and R.C.

## Data and materials availability

The datasets generated and analyzed during the current study are publicly available in the Gene Expression Omnibus: accession number GSE126799 (RNA-seq) and XXX (ATAC-seq)

---

**Fig. S1.**
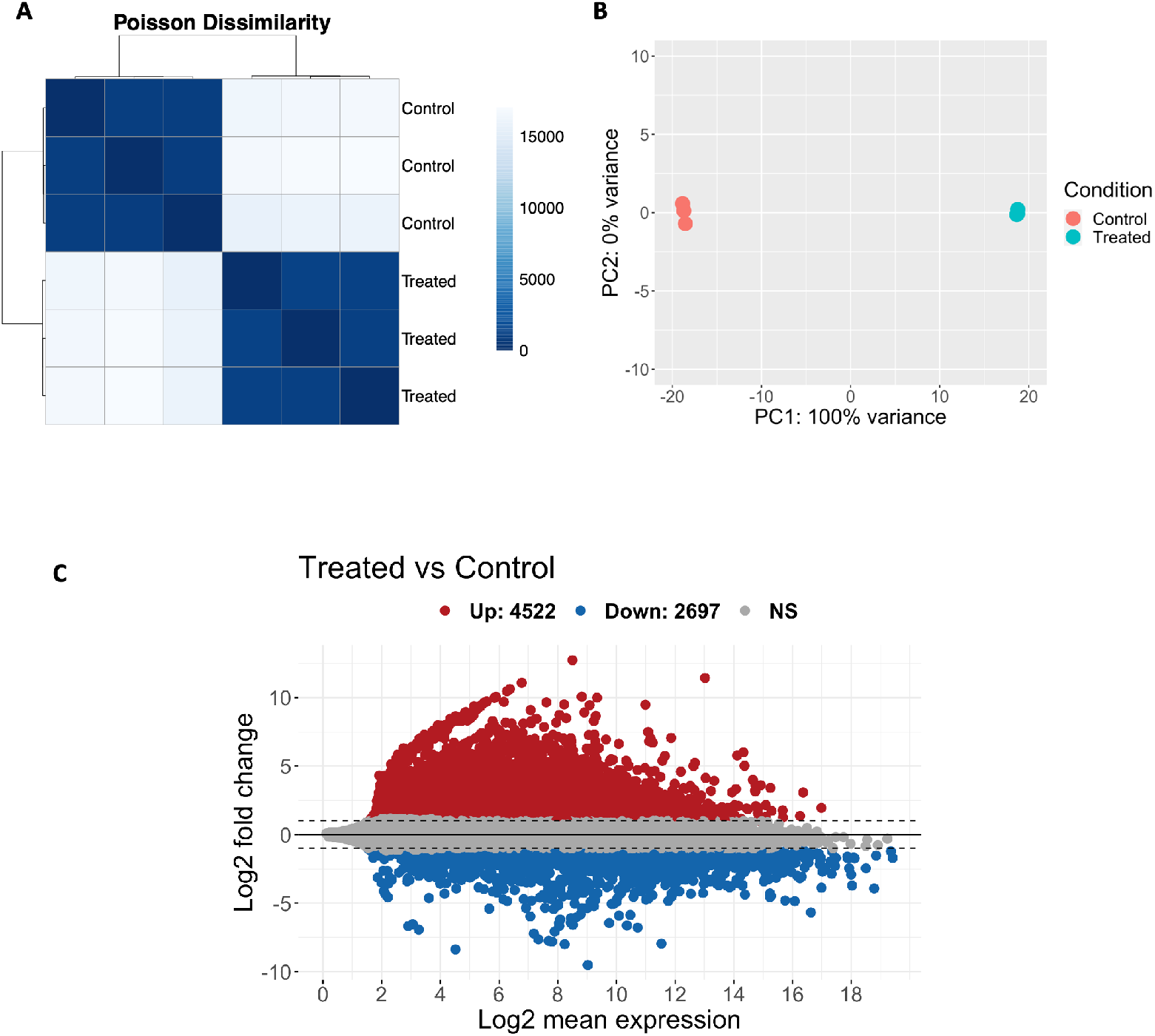
(A)Poisson distance clustering to overview the distribution of counts and clustering in the treated versus untreated group. Scalebar represents Poisson distance between samples. (B) Principal component analysis (PCA) of the DESeq2 analysis showing two distinct populations of control and treated group. PCA dimensionality reduction was performed on all samples. Almost 100% of the variance is associated with the first principle component, which separates replicates in the vehicle and octanoate treatment conditions. (C) DESeq2 analysis showing 2131 upregulated genes and 632 downregulated genes for octanoate group compared with the vehicle with FDR cutoff = 0.01 and |log_2_ FC| ≥ 2.

**Fig. S2.**
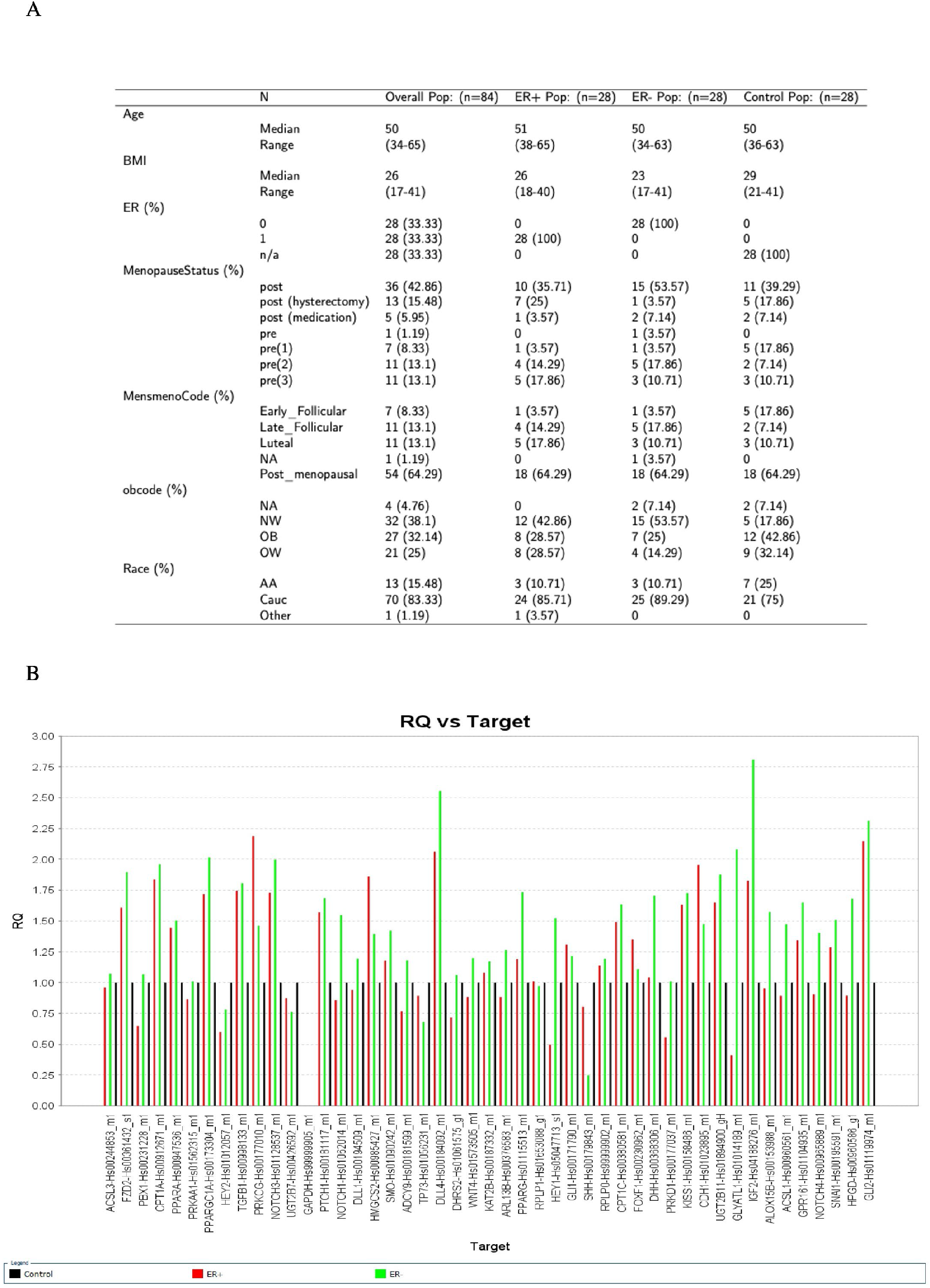
(A) The difference in age and BMI among three groups was analyzed by ANOVA with Sidak adjustment on pairwise comparison. The difference in menopausal status and race among three groups were analyzed using X^2^ test. The difference in HER2 status between ER1 and ER– group was analyzed using X^2^ test. (B) Histogram showing fold change or relative quantitation (RQ) for all genes of interest in the ER + (red) and ER- (green) in reference to the controls (black).

**Figure S3:**
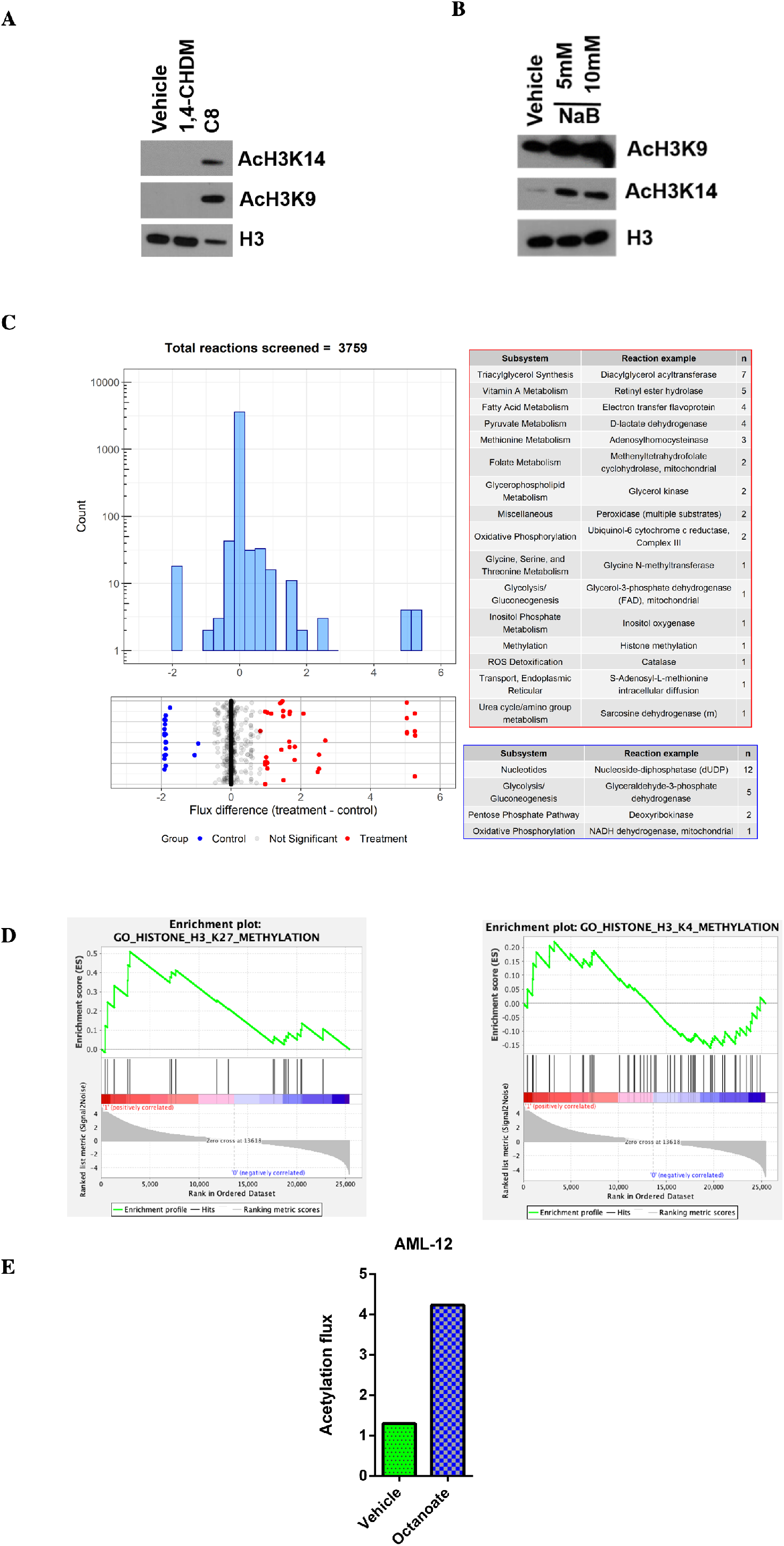
(A) To verify that the acetylation was specific to exposure to a fatty acid, MCF10A cells were exposed to 1,4-Cyclohexanedimethanol (1,4-CHDM), an alcohol with the same number of carbons, hydrogens and oxygens as octanoic acid. (B) Western blot of MCF-10A cells treated with HDAC inhibitor-sodium butyrate (NaB) 10mM for 24 hours to validate the specificity of the histone antibodies against the acetylated H3K9 and HK14. (C) The histogram and scatter plot show the distribution of flux differences of all 3759 metabolic reactions in the model between octanoate treatment and control. The horizontal x-axis shows the difference in flux of each reaction, while the y-axis of the histogram shows the total number of reactions in each bin. Metabolic pathways and representative reactions that showed the greatest differences in flux (p-value < 0.01) between the treatment and control are highlighted in the scatter plot and listed in the table. (D) GSEA analysis showing H3K27 and H3K4 enrichment in octanoate treated cells with corresponding leading-edge genes. (E) Predicted acetylation flux in octanoate treated AML-12 cells using FBA model.

